# Designing signaling environments to steer transcriptional diversity in neural progenitor cell populations

**DOI:** 10.1101/2019.12.30.890087

**Authors:** Sisi Chen, Jong H. Park, Jialong Jiang, Tiffany Tsou, Paul Rivaud, Matt Thomson

## Abstract

Stem and progenitor populations within developing embryos are diverse, composed of different subpopulations of precursor cells with varying developmental potential. How these different subpopulations are coordinately regulated by their signaling environments is not well understood. In this paper we develop a framework for controlling progenitor population structure in cell culture using high-throughput single cell mRNA-seq and computational analysis. We find that the natural transcriptional diversity of neural stem cell populations from the developing mouse brain collapses during in vitro culture. Cell populations are depleted of committed neuroblast progenitors and become dominated by a single pre-astrocytic cell population. By analyzing the response of neural stem cell populations to forty combinatorial signaling conditions, we demonstrate that signaling environments can restructure cell populations by modulating the relative abundance of pre-astrocytic and pre-neuronal subpopulations according to a simple log-linear model. Our work demonstrates that single-cell RNA-seq can be applied to learn how to modulate the diversity of stem cell populations, providing a new strategy for population-level stem cell control.

**Highlights:**

- Natural progenitor diversity in the brain collapses during in vitro culture to a single progenitor type
- Loss of progenitor diversity alters fate potential of cells during differentiation
- Large scale single-cell signaling screen identifies signals that reshape population structure towards neuronal cell types
- Signals regulate population structure according to a simple log-linear model

**GRAPHICAL ABSTRACT:** 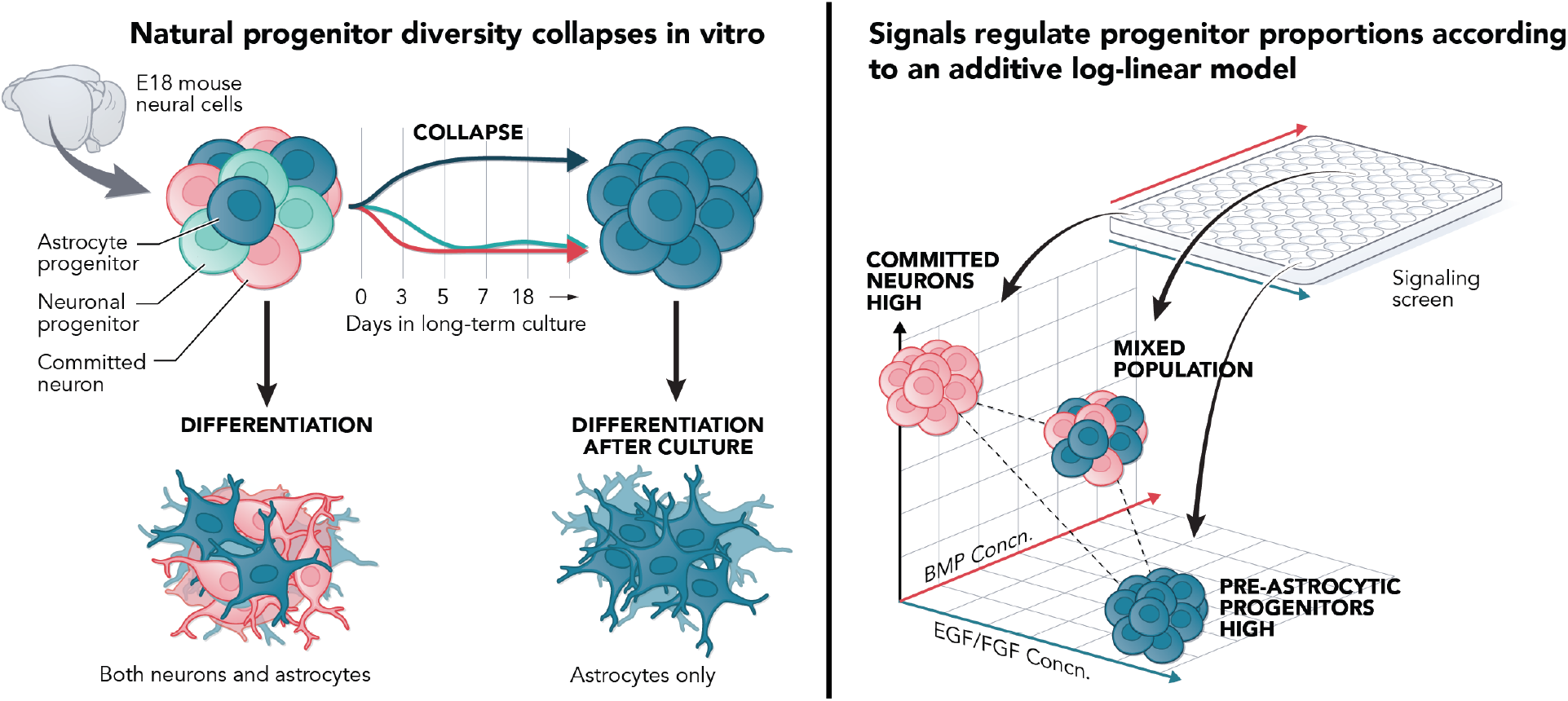

## Introduction

Stem cells are an exciting resource for regenerative medicine and tissue replacement. Work over many decades has shown that a wide range of stem cell populations, including embryonic stem cells[1] and neural stem cells [2], can be extracted from an embryo and then expanded and differentiated in cell culture using applied signaling molecules [3, 4, 5]. Expansion of cells in culture provides a strategy for generating a wide range of functional cell types from a small initial pool of precursors that could be used to replace or repair damaged tissue in the brain[6], heart[7], and other organs that do not have intrinsic regenerative capacity.

However, despite the promise, the control of stem cell populations in cell culture, outside of the embryonic environment, has continued to be difficult and limited in many cases. Many cell types of interest are still not conducive to induction and growth in vitro. For instance, the striatal medium spiny neurons that are central to Huntington’s disease cannot be generated with sufficient functional properties or high enough efficiencies to be useful as a model of the disease[8]. As another example, mature T-cells cannot be generated from pluripotent stem cells without the presence of a complex artificial niche architecture [9] and even then, do not replicate certain key features of T-cell phenotype [10, 11]. The differentiation process can also be highly variable and a poor understanding of the nature and source of this variability has led to practical bottlenecks in research and costly failures in clinical trials [12, 13, 14].

Part of the challenge to manipulating stem and progenitor cells has been a poor understanding of the heterogeneity of the populations we can manipulate in culture. During development, the embryo regulates distinct pools of multipotent stem and progenitor cells [15, 16, 17] to generate a diversity of mature cell types in the correct proportions. Changes in the population composition of lineage-restricted precursors [18, 19] can dictate the relative abundance of neurons versus astrocytes during brain development, or, in the immune system, the relative abundance of lymphoid versus myeloid cells in disease or aging [20]. Altering the progenitor cell distribution has also been shown to impact downstream differentiation during the differentiation of induced pluripotent stem cells [21]. However, despite the importance of stem and progenitor cell population diversity in the developing embryo, we typically do not know how natural progenitor diversity changes within cell culture signaling environments or how to shape population diversity to amplify progenitor types of interest.

In this work, we demonstrate a new approach to programming stem cell populations by using combinatorial signaling ligands to directly steer the population composition of precursor cells isolated from the embryonic mouse brain. Here, we apply single cell mRNA-seq to resolve distinct progenitor substates from both the developing mouse brain and cells that are extracted and grown in culture (SI Table S1). In culture under canonical EGF/FGF signaling conditions, pre-neuronal progenitor states are lost, leaving only progenitor cells that primarily generate astrocytes upon differentiation. By exposing dissociated cells to forty different signaling environments, we show that the signaling environment can alter the balance of stem cell populations and amplify pre-neuronal populations that are lost under canonical conditions. Moreover, we found we could model the population response using a simple log-linear model, suggesting that most of the signals we assayed have effects on proliferation that combine additively. Our work points to population structure and dynamics as playing an important and previously under-appreciated role in impacting the outcome of stem cell differentiation. Broadly, we demonstrate an experimental and computational strategy for designing signaling environments that can restructure cell populations to overcome challenges implicit in stem cell therapies.

## Results

### Neural stem cell populations experience collapse in transcriptional diversity in cell culture

We wanted to understand how the diversity of neural stem cell populations changes when cells are extracted from the embryonic brain and placed in cell culture signaling environments. Therefore, we applied single cell mRNA-seq to create a map of the stem cell populations contained in the developing mouse brain, and then compared the underlying transcriptional diversity to neural stem cells grown in cell culture. Specifically, we profiled 33,998 cells at three time points of embryonic development (embryonic day 18, postnatal day 4, and Adult Cortex) and compared these cells to populations of stem cells dissociated from the E18 brain and cultured as neurospheres using canonical growth factors, EGF2 and FGF2 (Figure 1a). From the E18 brain, we specifically dissociated a subportion of the brain containing the subventricular zone, which harbors the stem and progenitor cells that give rise to the entire cortex.

**Figure 1:**
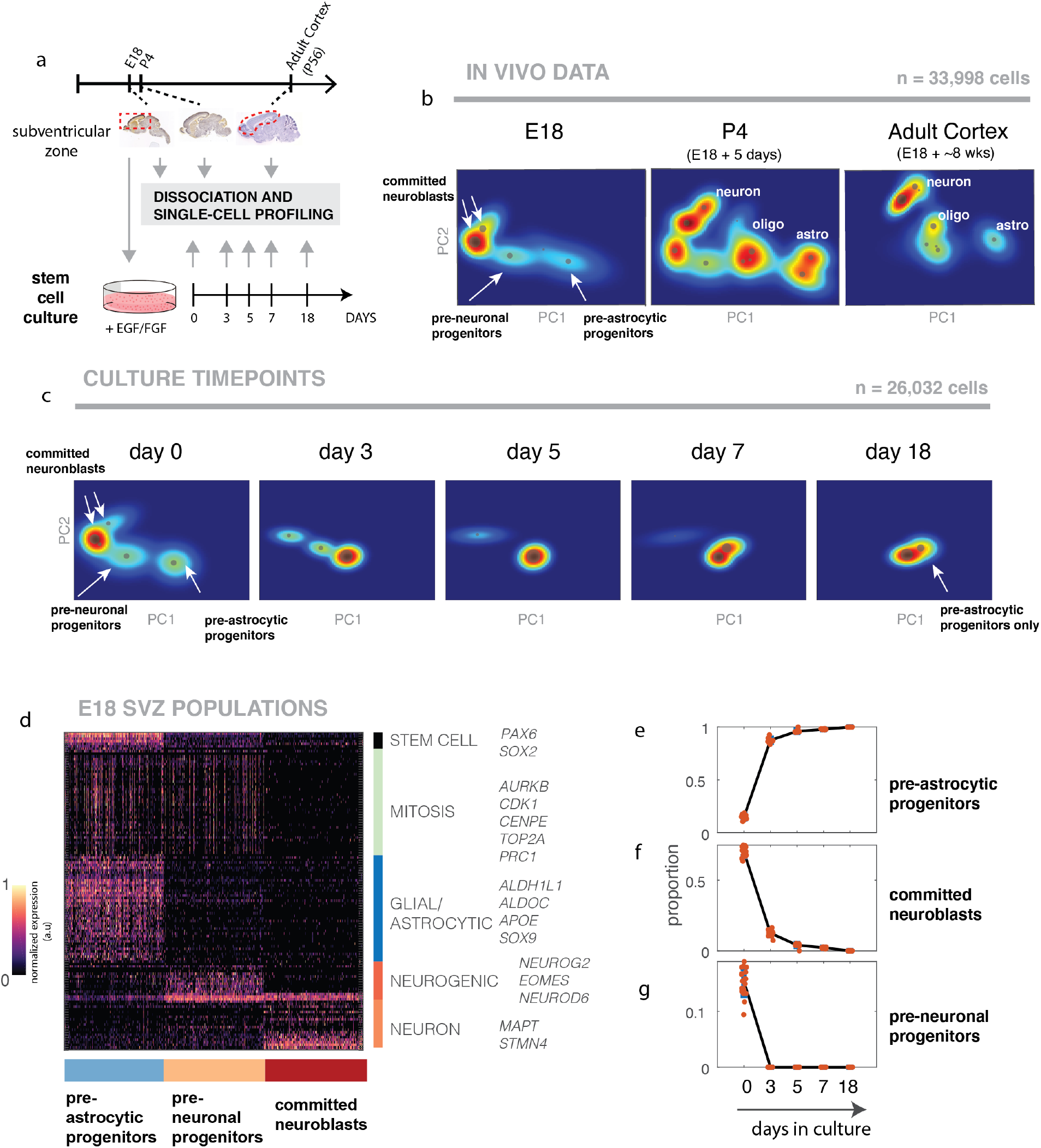
Transcriptional diversity of embryonic neural stem cell populations collapses in vitro. (a) Experimental scheme showing timepoints at which cells are profiled from the brain (upper) and during stem cell culture using canonical growth factors (EGF/FGF2). Stem cell cultures are initiated from cells dissociated from the E18 subventricular zone. (b) 2D-projected renderings of GMM probabilistic model built for each dataset collected directly from E18, P4, and Adult cortex tissues. Each GMM describes a probability distribution of cell populations within a 10D PCA space. The mean gene expression state of each independent Gaussian is displayed as a dot scaled by population abundance. (c) 2D-projected renderings of GMM probabilistic model built for each timepoint during a culture timecourse. The renderings show a qualitative collapse of diverse progenitor populations to only the pre-astrocytic progenitor population. (d) Gene expression heatmap of the three identified cell subpopulations from E18 brain. Each cell type is composed of expression of one of six indicated gene expression programs. Pre-astrocytic progenitor cells expresses stem cell genes, mitotic genes, as well as a glial/astrocytic gene program. Pre-neuronal progenitor cells expresses stem cell genes, mitotic genes, as well as a neurogenic gene program. Committed neuroblasts express neuronal genes, but not the other programs. (e-g) Quantification of proportions across timecourse for each major subpopulation type (e) pre-astrocytic progenitors, (f) Committed neuroblasts and (g) pre-neuronal progenitors. Proportion values are bootstrapped over multiple model building runs to average out potential variability in model fitting. For each timepoint, we built 20 distinct models (red points), and aligned subpopulations to reference subpopulations (day 0 timepoint) to associate abundance values.

To analyze the data, we used a probabilistic modeling platform, PopAlign, [22] to build, align, and compare Gaussian mixture models (GMMs) (SI Figure S1) of the embryonic (Figure 1b) and cell culture populations (Figure 1c). Cell type markers for mature differentiated cell types (SI Figure S2a) are used to type subpopulations in the postnatal day 4 and adult cortex samples (SI Figure S2b), which provide a reference for assessing differentiation potential. For each sample, the resulting PopAlign models were low error both qualitatively (SI Figure S3) and quantitatively (SI Figure S4) and so provided a computational substrate for performing quantitative comparisons between cells directly extracted from tissue and their cultured derivatives.

Our computational analysis revealed that embryonic day 18 (E18) brain normally contains a broad range of pre-neuronal and pre-astrocytic progenitor cells (Figure 1b) but that this population diversity collapses over time in cell culture (Figure 1c). Specifically, in the E18 subventricular zone, we identified three major categories of cells as the centroids of the GMMs identified by PopAlign (Figure 1b, 1c): committed neuroblasts, pre-neuronal progenitor cells, and pre-astrocytic progenitor cells. We labeled these distinct progenitor populations based on their expression patterns of lineage-specific and mitotic genes expression programs (Figure 1d). Both pre-neuronal and pre-astrocytic progenitor populations expressed genes associated with lineage-specific fate (neuronal: Neurog2; astrocytic: Sox9), while still expressing stem cell transcription factors (Pax6, Sox2) and mitotic genes. Committed neuroblasts expressed a neuron-specific gene expression program, but did not express stem cell genes or the mitotic gene expression program.

Long-term culture with canonical growth factors, EGF and FGF, caused the diversity of this entire population to dramatically shift, undergoing a qualitative loss of the pre-neuronal and committed neuroblast subpopulations within five days of culture (Figure 1c). Initially, committed neuroblasts comprise 71.4% of the population, while pre-neuronal progenitors and pre-astrocytic progenitors make up the rest at roughly even proportions (13.8% and 12.4% respectively). By day five of culture, the cell population had become dominated by the pre-astrocytic progenitor cells (96% of cells at day 5 but 18% at day 0) (Figure 1e) and the committed neuroblast cells have been lost (4% day 5 vs 66% day 0) (Figure 1f). Furthermore, pre-neuronal progenitors diminish from 16% to 0% at day 5 (Figure 1g). In this way, although the developing brain at E18 contains a diverse population of progenitor types, only the pre-astrocytic progenitors are expanded in EGF/FGF2 cell culture.

Previous work on neural stem cells extracted from the mammalian brain have suggested that cultured neurospheres maintain transcriptionally heterogeneous populations [23]. However, our results demonstrate the opposite - that these canonical factors actually trigger a collapse to the pre-astrocytic progenitor state (Figure 1c), which occurs dynamically over the course of multiple days. Although many reasons could give rise to this discrepancy - a difference in the underlying species used (human vs mouse) and the dynamics of population collapse (Short-term vs long-term cultures) - we note that previous methods could not resolve the difference between gene expression stochasticity and the presence of distinct subpopulations.

### Decreased population diversity is associated with limited differentiation potential

Next, we investigated the impact of population collapse on the types of differentiated cells that can be generated, finding that the collapsed population of pre-astrocytic progenitors had restricted fate potential.

We differentiated cells immediately after dissociation from the E18 subventricular zone, as well as cells from the collapsed population that had been cultured in EGF and FGF for 7 days. Upon differentiation in serum-containing media, the EGF/FGF cultured cells generated uniform cultures of astrocytes with no density in the region corresponding to the neuronal branch of the cell population at postnatal day 4 (P4) (Figure 2b), which temporally corresponds to 5 days of in vitro differentiation. These differentiated populations quantitatively align most closely to the P4 astrocytic population, and produce matching gene expression signatures (Figure 2c).

**Figure 2:**
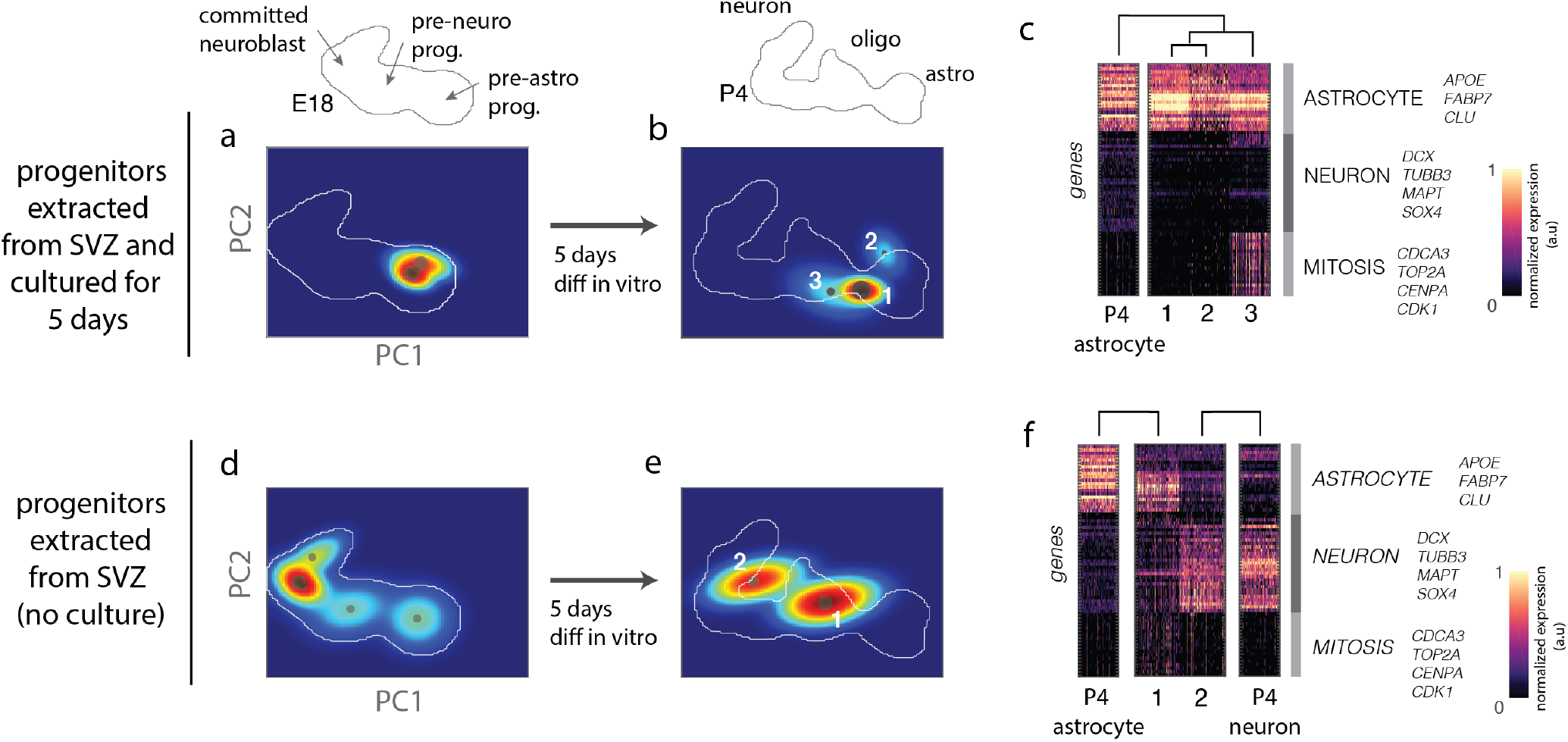
Decreased progenitor diversity is associated with limited differentiation potential. (a) PCA rendering of cell populations extracted from the SVZ and cultured for 5 days in EGF/FGF and (b) after differentiating in serum-containing NbCo-Culture media for 5 days. Above: map outline of PCA manifolds in the reference E18 SVZ sample or the P4 sample, with cell states labeled for reference. (c) Gene expression heatmaps of population centroids from differentiated cells, compared to aligned populations from the in vivo brain at P4. Astrocytic, mitotic, and neuronal gene expression programs are indicated in the key and supplied in Supplementary File 2. (d) Model renderings of cells dissociated directly from the SVZ and (e) their differentiated progeny after 5 days of in vitro differentiation. (i) Gene expression heat-map comparing the centroids of populations from (e) to their aligned in vivo populations.

However, under the same conditions, cells extracted from SVZ tissue (Figure 2d) and immediately differentiated could generate mixed cultures of neurons and astrocytes based on their locations within a globally defined PC projection (Figure 2e), and gene expression profiles (Figure 2f). Although we cannot definitively conclude that the pre-astrocytic progenitor cells lack neuronal potential under all possible contexts, our results show that they do not generate neurons in conditions that typically support neuronal differentiation (Serum supplementation and EGF/FGF withdrawal). The pre-astrocytic nature of stem cells is likely due to the fact that the cells were isolated at E18, when the neural stem cells in the fetal cortex are undergoing the gliogenic switch[24]. Thus, these findings suggest that the collapse in progenitor cell diversity during cell culture limits the capacity of the population to generate neurons, and imply that increasing the progenitor diversity could expand the ability to produce neuronal cell types.

### Combinatorial signaling environments modulate balance between pre-astrocytic progenitors and committed neuroblasts

We then wanted to understand whether it was possible to recover or manipulating progenitor cell type diversity, which could be important for altering fate potential.

Canonical neural stem/progenitor culture involves the use of EGF and FGF signaling ligands [25, 26, 27, 28, 29, 30, 31, 32, 33] to stimulate stem and progenitor cell proliferation. In our experiments, the EGF and FGF2 used in canonical neural stem cell culture selectively amplify only the pre-astrocytic progenitor cells, which might thus restrict the differentiation potential of the cell population. This population selection happens at the expense of the pre-neuronal progenitors and committed neuroblasts, whose proportions decline over long-term culture in EGF/FGF. We note that this difference is not due to the absence of EGF and FGF receptors on pre-neuronal progenitors and committed neuroblasts. Both of these populations express the receptor for FGFR1, one of the main receptors for FGF2 (SI Figure S5). Additionally, FGFR2, FGFR3, and EGFR are also expressed to some extent in the pre-neuronal progenitors (SI Figure S5).

While EGF and FGF2 at a fixed concentration represent just a single signaling condition, the embryonic brain contains combinatorial gradients of many distinct signaling pathways [34, 35]. Therefore, we asked whether we could stabilize a more diverse progenitor population by employing additional signaling factors. We performed a signaling screen where we exposed the SVZ stem cell populations across a range of different signaling conditions and analyzed the impact of signaling combinations on population structure using single-cell transcriptional profiling.

We selected signaling cues from the EGF, FGF, Wnt, PDGF, and BMP families whose receptors are all present in the developing brain (SI Figure S5) and are known to be associated with diverse functions such as proliferation (EGF,FGF2 [36], neuronal differentiation (WNT[37] PDGF), astrocytic differentiation (BMP4[38] and FGF9[39]), and the maintenance of quiescence (BMP4 [40]). We also included two immune signaling cytokines, GM-CSF and IFN-gamma as outgroups that are not normally active during neural development, but whose receptors are present in these cells (SI Figure S5).

We combined all signals with EGF/FGF2 in varying doses ranging from 20ng/mL, a high concentration that is the standard culture condition, down to 0.8 ng/mL, a concentration that we found was necessary to maintain cell survival (SI Table S2). Signals were added immediately to cells after SVZ dissociation. Cells were profiled after 5 days of exposure, and the Clicktags multiplexing platform [41] was applied to label individual samples. We profiled an average of 500 cells per signaling condition leading to a data set with 18,700 cells, and applied PopAlign to dissect and analyze population diversity and to compare identified cell populations to those in the E18 brain.

We found that signaling conditions could shift the balance between pre-neuronal and pre-astrocyte cell populations. Across all signaling conditions, we identified three major subpopulations of progenitor cells that aligned to the three cell populations we previously identified (Figure 1) in the E18 mouse brain: committed neuroblasts, pre-astrocytic progenitors and pre-neuronal progenitors (Figure 3a, SI Figure S6). Individual signals altered the relative abundances of the pre-neuronal and pre-astrocyte subpopulations, with the subset of conditions including BMP showing particularly strong impact. Specifically, BMP4 shifted the population structure towards more pre-neuronal cells (committed neuroblast - pink), which could be observed qualitatively in terms of increasing density in the committed neuroblast population (Figure 3b) and quantitatively in a phase-diagram (Figure 3c). Increasing the concentration of EGF/FGF2 had the opposite effect, shifting the population away from committed neuroblasts, towards pre-astrocytic progenitors (Figure 3b, 3d). CHIR boosted the proportions of pre-astrocytic progenitors (SI Figure S7a), while having no effect on committed neuroblasts (SI Figure S7b). Other signals such as FGF9, PDGF-AA, GM-CSF, and IFNG predominantly reduced the proportions of pre-astrocytic progenitors (SI Figure S7c) and boosted committed neuroblasts (SI Figure S7c).

**Figure 3:**
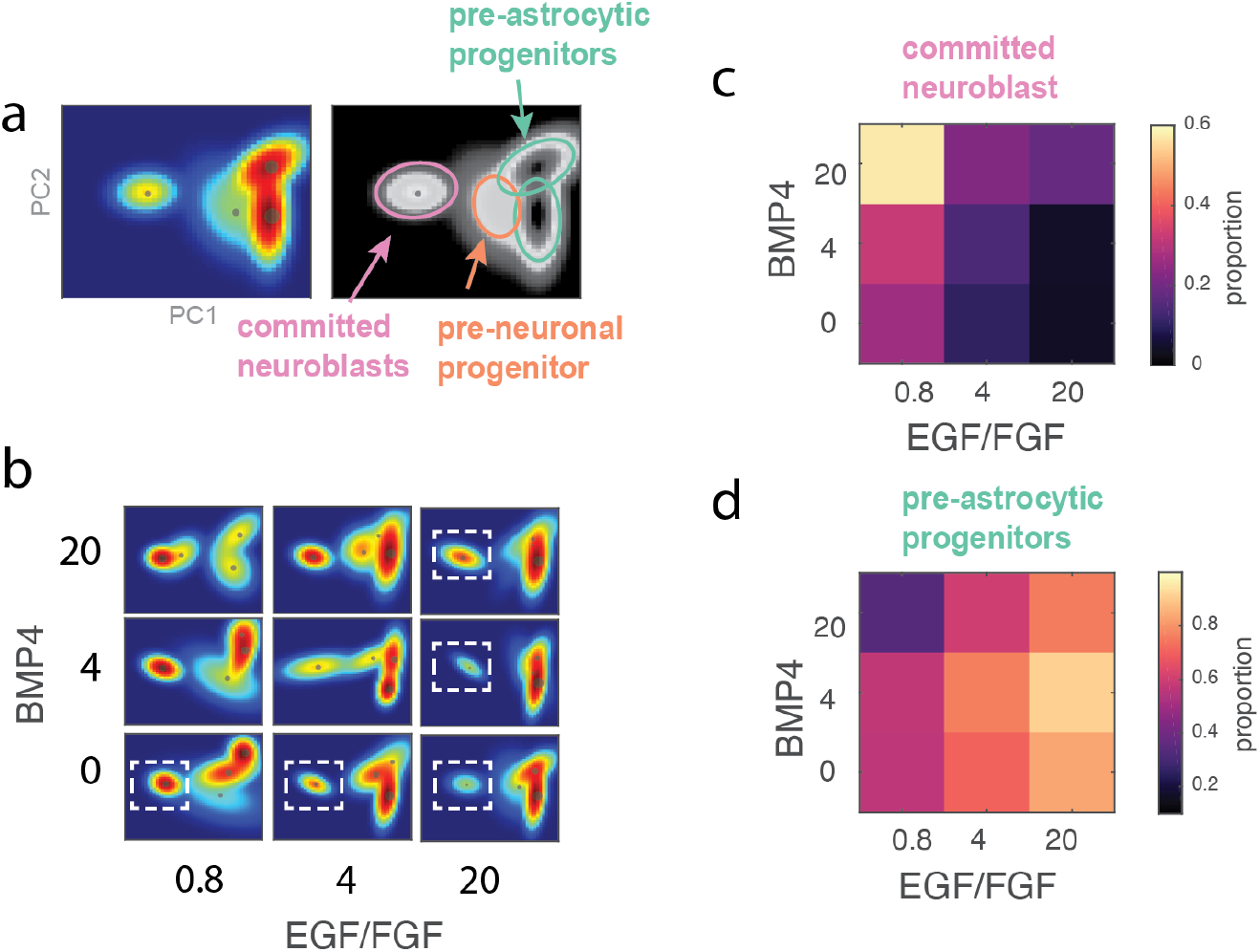
Signals reshape global population diversity. (a) 2D PCA rendering of common model built using single-cell data sampled across 40-condition signaling screen Identified subpopulations align to the main subtypes from the in vivo SVZ-D0 population: pre-astocytic progenitor, committed neuroblast, pre-neuronal progenitor. (b) Renderings show population shifts in response to varying EGF/FGF and BMP4. The density of the committed neuroblast population, highlighted in dashed white box, increases with increasing BMP4, and decreases with increasing EGF/FGF. (c) Heatmap of committed neuroblast proportions with varying concentrations of BMP4 and EGF/FGF. Increasing BMP4 increases the committed neuroblast proportion, while increasing EGF diminishes it. (d) Heatmap of pre-astrocytic progenitor proportions, showing it increasing with EGF/FGF, and decreasing with BMP4.

### Simple log-linear model enables design of signaling environments to achieve target population structure

The combinatorial impact of the signaling environment on the relative abundance of the three underlying progenitor cell populations could be captured by a simple log-linear model. To capture the relationship between signaling inputs and population size, we consider a simple log-linear model where the relative growth rates of specific cell populations respond independently to signaling inputs (Figure 4a). In this model, the log of the relative proportion of population (e.g. committed neuroblast/pre-neuronal progenitor) depends on a linear combination of signaling molecule concentrations (Figure 4b). We use the relative proportion of subpopulation *i* with respect to a reference subpopulation *k* to deal with the loss of one degree of freedom since the total population proportions sum to 1 (See SI Methods). The relative proportions are computed directly by using the PopAlign abundance parameters *w*_*i*_/*w*_*k*_.

**Figure 4:**
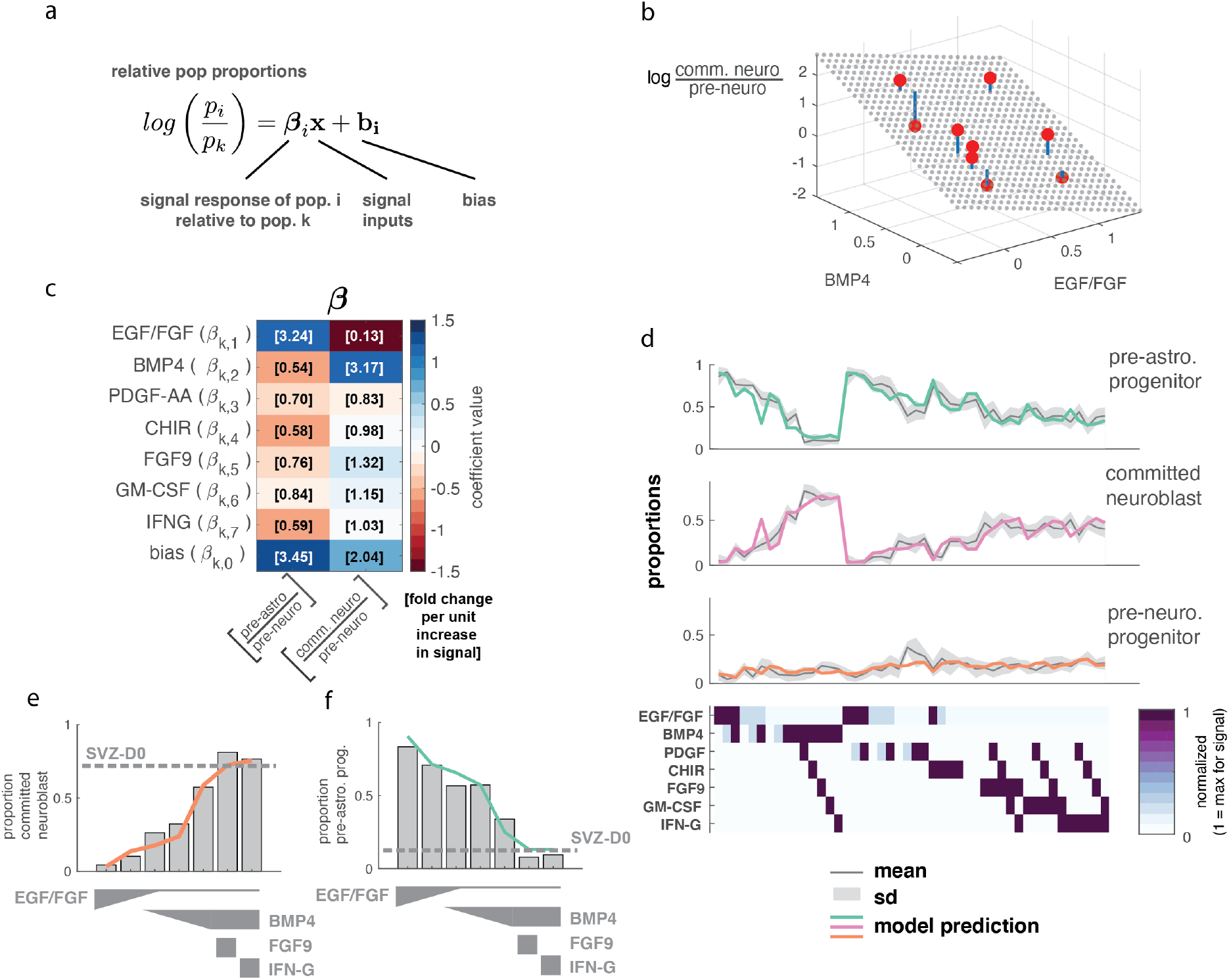
Population structure responds to signaling input according to a simple log-linear model. (a) Model equation relating relative subpopulation proportions (*p*_*i*_*/p*_*k*_) to a linear function of signaling inputs, Coefficients *β*_*i*_ can be considered the relative growth rate of population *i* to population *k*. The bias encodes the natural difference in growth rates in the absence of signal (***x*** = 0). (b) The relative proportion of committed neuroblast to pre-astrocytic progenitor populations (logged) responds linearly to signaling inputs. Grey dots: model output, red dots: observed proportions, blue line: error between observed data and prediction. (c) Coefficient matrix *β* linking relative population ratio (columns) to signals (rows). The number in brackets denotes the exponent of the coefficient value 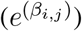, and can be interpreted as the fold change in relative proportions *p*_*i*_*/p*_*k*_ caused by each unit increase in signal. (d) Observed and predicted (colored) subpopulation proportions reported for each experimental sample (column). The standard deviation is calculated by bootstrapping samples from experimental data (n=100 cells each, iterations=20). Heatmap represents signaling concentration used within the column. (e-f) Model (colored lines) predicts combinations of signals that push committed neuroblast and pre-astrocytic progenitor proportions towards target levels, such as that seen in original tissue (dotted line - SVZ-D0). Note: Signal concentrations across all panels in this figure is normalized so that 1 unit equals the maximum concentration (all 20 ng/mL except IFN-G and GM-CSF, whose max concentration is 3 ng/mL, and CHIR, whose max concentration is 3 uM).

The coefficients *β* of the log-linear model allow us to interpret the effects of each signal on a specific subpopulation. Each entry *β*_*ij*_ can be interpreted as the difference in relative growth rate between subpopulation *i* versus reference subpopulation *k* for each signal *j*. Importantly, *β*_*ij*_ can encode both positive and negative values and can reveal antagonistic relationships between signals (Figure 4c). For instance, BMP4 and EGF compete to modulate the relative population sizes of both committed neuroblast and pre-astrocytic progenitor cells (relative to the pre-neuronal progenitor population, which we take to be reference population *k*) (Figure 4c). Model parameters reveal that each additional unit of BMP4 increases the relative proportion of committed neuroblasts by 4.4x, while EGF/FGF reverses this trend, scaling down the relative committed neuroblast proportion by 0.4x (Figure 4c).

The simple log-linear model was sufficient to recreate observed population abundances across all experiments. The accuracy of the model is both qualitative and quantitative. Model-generated predictions match patterns seen in the measured subpopulation abundances (Figure 4d). Additionally, the mean absolute error (MAE) between the model-predicted proportions and the observed proportions is small, averaging only 6.52% of the total population. The accuracy of the simple log-linear model suggests that additive impact of signaling molecules on cell-type specific proliferation rates might be sufficient to explain much of the population restructuring observed across the combinatorial signaling conditions assayed in our screen (SI Methods).

Using the idea of a linear signaling code, we were able to design combinatorial signaling conditions to achieve desired population structures. For instance, although none of the signaling combination we tested could completely recreate the transcriptional diversity of the SVZ tissue, some conditions could approximate it by suppressing unwanted pre-astrocytic populations and boosting desired pre-neuronal populations. The original SVZ tissue was composed of *>* 70% committed neuroblasts (Figure 4e-dotted line). By reducing EGF/FGF2 alone, we boosted the population of committed neuroblasts to 26% of the population (Figure 4e). Adding BMP4 boosts the population percentage to 56%. Finally, adding another committed neuroblast-boosting and pre-astrocytic progenitor-reducing signal such as FGF9 or IFNG can boost the committed neuroblasts again up to *>* 80%, while also reducing the number of pre-astrocytic progenitors to 4-6% (Figure 4g). Globally, the conditions satisfying these constraints (low EGF/FGF2, high BMP4, and an additional pre-astrocytic progenitor-reducing signal) produced populations of cells that closely match the original SVZ tissue at day 0, based on a statistical metric which calculates the divergence[22] of the two models (SI Figure S8).

## Discussion

The ability to manipulate the relative proportions of distinct stem cell and progenitor subpopulations would provide an important route to controlling the differentiation potential of a heterogeneous stem cell population and also to modulating the state of tissues in the body during development. In our work, we apply single-cell profiling and large-scale combinatorial signaling experiments to dissect how individual signals and combinations of signals alter the population structure of progenitor cells extracted from the developing E18 mouse brain. We showed that under canonical culture conditions, progenitor cells naturally collapse into a single pre-astrocytic progenitor population, but that we can reshape the population proportions by modulating the concentrations of a set of signals found in the developing brain (EGF, FGF2, FGF9, Wnt, BMP, IFNG, and GM-CSF). We further show that these signaling proteins impact populations proportions in a way that can be modeled by a simple log-linear model, suggesting additivity in the way that their effects combine.

Our approach has implications for understanding the behavior of stem cell populations broadly. Developing cells undergoing morphogenesis in the embryo are exposed to combinations of signaling molecules[42] generated by morphogen gradients[43, 44, 45]. An important question is to determine how stem cells can decode combinatorial signaling environments to select their fate and proliferation rate. Our findings point towards a potentially simple signaling code in which different signals linearly regulate the proliferation rate of distinct transcriptionally primed progenitor cell populations. By systematically expanding upon this code across a wider ranger of signals, we can develop strategies for selectively growing and shrinking different populations of progenitors within a heterogeneous culture, which would be a new strategy for directing stem cell fate for therapeutic applications.

Our result that the population responses to signals can be modelled with an additive model may seem surprising in light of other research suggesting that combinations of signaling ligands have more complex properties [42, 46, 47, 48]. Development has many examples of a combinatorial signaling code generating specific fates, such as the generation of mesodermal cells [46] or the precise patterning of individual cells within the fly eye ommatidia [47]. More recently, Antebi et al. showed that pairwise combinations of BMP ligands yield many different categories of response, beyond additive responses, including imbalance detection (which behaves like an XOR gate), balance detection (which behaves like an AND gate), or ratiometric response, in which the response scales according to the ratio of two ligands[48].

However, multiple reasons could explain why an additive log-linear model is sufficient for describing the combinatorial signaling responses we have observed. First, additivity in biological signal processing is a general principle that appears to be the default mode of computation. Antebi et al. show even show results suggesting that additivity is the most common type of signal integration in an *in silico* ‘screen’ of pairwise interactions of simulated ligands. Second, the particular signals that we have tested are from distinct ligand families, whereas the signals in [48] are from the same family of BMP ligands. It is well known that structurally similar proteins (i.e. that emerge from gene duplication events[49, 50]) can evolve altered binding properties to confer new types of activity, such as receptor competition [50, 51]. Thus, we conjecture that interactions between ligands of different families may be less likely to exhibit complex non-additive behaviors, like antagonism or imbalance detection. Third, the combinatorial impact of a signal depends on the nature of the response variable. While most combinatorial response studies have been focused on regulation of single genes or reporters, in our study we considered the impact on composition of an entire population of cells, which is an integrated response across many layers of regulation (from signal transduction to transcriptional regulation of proliferation) as well as across many different cell types. These layers of regulation may potentially buffer any nonlinearities in the acute response to signals. Finally, we emphasize that fuller exploration of the combinatorial signaling space will allow us to uncover interesting non-linear regimes, but will involve measuring thousands of signaling conditions across doses and combinations. Single-cell RNA-sequencing experiments of this scale are still largely cost-prohibitive but are becoming increasingly feasible with new technologies [41, 52, 53, 54].

Our work emphasizes the role of local signaling environments in generating and maintaining population diversity both in the embryo and in artificial cell culture contexts. The embryo maintains population diversity naturally, by controlling signaling environments on cellular length scales to allow divergent cell-types to be generated in close spatial proximity [55]. The population diversity of the embryo is thus directly enabled by the self-organization of local combinatorial signaling environments. An important challenge for cell engineering is to control combinatorial signaling environments outside of the body with similar precision. In engineering applications, signals have traditionally been applied globally, to an entire cell culture, and therefore, might be fundamentally limited in their ability to control the generation of different cell types in close spatial proximity. Organoid-based approaches solve the signaling control problem by generating signals locally through cell-cell interactions [56]. Future and current engineering approaches [57, 58] in which signals can be modulated on cellular length scales will be critical for learning how different cell-types can emerge through distinct, micron-scale, local signaling environments.

Broadly, our approach can be applied to study the combinatorial impact of a wide range of signaling molecules on heterogeneous populations of stem and progenitor cells. The key technical innovations we apply are single-cell mRNA-seq, population level gene expression analysis and computational modeling. We anticipate that this strategy can be applied to a much broader range of signals to uncover the underlying signaling code for engineering the population structure of heterogeneous multicellular constructs used in both synthetic biology and tissue engineering applications.

## Methods

### Mouse brain tissue dissociation

Tissue from mouse brain was purchased from Brainbits or acquired from collaborators (Allan Pool, Oka Lab, Caltech, Elisha McKay, Gradinaru Lab), and digested using 6mg of Papain (Brainbits) at 2mg/mL in Hibernate E medium(Brainbits) for 30 minutes at 30°C. After incubation, tissue was triturated for 1 minute using a silzanized pasteur pipet. Dissociation medium containing the disociated cells were centrifuged for 1 minute at 200 × g and the cell pellet was either resuspended in media for further culture or resuspended in 1x PBS + 0.04% BSA for single-cell profiling through the 10X Genomics platform.

### SVZ cell derivation

Neural progenitors are dissociated from the combined cortex, hippocampus and ventricular zone of E18 mice (Brainbits). After dissociation and resuspension of the cells in NPgrow media (Brain-bits), cells were counted and plated onto ultra-low attachment 100mm culture plates (Corning) at one million cells per plate in 10mL of NPgrow, and supplemented with 1% Penicillin-Streptomycin (Gibco). NPgrow media consists of Neurobasal media, 2% B27 (minus Vitamin A), 0.5mM Glutamax, 20 ng/mL EGF and 20ng/mL FGF. Cells were cultured at 37°C and 5% CO2 and supplemented medium was changed every 2 to 3 days.

### Differentiation

In vitro-expanded and directly dissociated progenitor cells were differentiated for 5 days in custom Brainbits NbCo-culture media that supports the growth of both neuronal and glial differentiated cells. Media consists of Neurobasal, 2% B27 (minus Vitamin A), 0.5mM Glutamax, 10% FBS, and 1% Penicillin-Streptomycin (Gibco), and was supplemented with a proprietary mixture of creatine, estradiol, and cholesterol. Cells were seeded onto poly-ornithine/laminin coated plates at a density of 50k cells/cm^2, and cultured at 37°C and 5% CO2. Medium was changed every 2 to 3 days. Cells were cultured for 5 days before dissociation for single-cell profiling.

### Combinatorial signaling screen

We applied combinatorial signals to cells dissociated from the SVZ region of the E18 mouse brain to understand how signals regulate the population structure and transcriptional state of progenitor cells. We used NPGrow media which did not contain EGF/FGF, and added signaling factors to the final concentrations given in SI Table S2. For EGF, FGF2, BMP4, PDGF-AA, and FGF9, the maximum concentration used was 20 ng/mL. For IFN-G and GM-CSF, the maximum concentration used was 3 ng/mL, and for CHIR99021, the maximum concentration used was 3 uM. After cells were resuspended in media, they were cultured for 5 days before dissociation for clicktag multiplexing and single-cell profiling.

### Experimental multiplexing using Clicktags

Cultured cells were dissociated in Accutase into a single cell suspension. Cells were then fixed in ice-cold 80% methanol. Prior to cell labeling, two methyltetrazine (MTZ) activated barcoding oligos were combined for each labeling sample, NHS-trans-cyclooctene (NHS-TCO) was then added and left to reaction for 5 minutes. The previously fixed cells were then mixed thoroughly with the barcode oligo solution and incubated for 30 minutes at room temperature. Labeling reaction was then quenched with the addition of Tris-HCl and methyltetrazine-DBCO. After the quenching step, cells from all labeling samples were pooled together along with the addition of two volumes of PBS-BSA then centrifuged at 800 x g for 5 minutes followed by two more cycles of PBS-BSA washes and centrifugation steps. After the final centrifugation step, the cell pellet was resuspended in 100uL of PBS-BSA and counted on a hemocytometer. Cells were then loaded onto 3-4 lanes, targeting 10,000 cells per lane, of the Chromium Controller and processed as recommended by the 10x Genomics v2 Single Cell 3’ reagent kit protocol until completion of the cDNA amplification PCR step. At this step, SPRI size-selection was used to separate the barcode oligos from the cDNA which were then processed separately. We used a dual-tagging approach, so that each experimental sample gets two clicktag barcodes.

### Barcode demultiplexing

Filtered cell barcodes from the aligned gene expression (Cellranger) is used as a whitelist to parse clicktags data from fastqs. Tag sequences extracted from fastqs are aligned to a list of known clicktag sequences, allowing for 1 mismatch (Levenshtein distance ¡=1). Counts for clicktags are accumulated and stored in a cell barcodes by tags matrix. To threshold the data, we computed a count distribution for each individual clicktag, and used Otsu’s method to obtain optimal thresholds for each clicktag sequence. Each cell is thresholded for each barcode independently, and cells which are have the correct barcode pair combinations are retained.

### Computational Methods

Detailed computational methods are given in the Supplementary Information. Briefly, single-cell data across all samples were analyzed using a Matlab implementation of the PopAlign framework to perform normalization, filtering, dimensionality reduction, Gaussian mixture model building, and subpopulation alignment [22]. To track subpopulations across samples, subpopulations in each sample were aligned to reference subpopulations found in the E18 sample. For multiplexed datasets, populations were analyzed using a Query function (See SI Methods - Query), in which a universal Gaussian mixture model is built from all of the data, and used to score cells as belonging to a particular subpopulation (i.e. independent Gaussian model). These class labels are used to compute proportion values *w*_*i*_ for each subpopulation. These values are converted into a relative abundance by dividing *w*_*i*_*/w*_*k*_, where *k* is the index of the selected reference population (pre-neuronal progenitors) within the dataset. The relative abundances are used as input into a log-linear model (Figure 4a, SI-Methods).

## Data Availability Statement

Single-cell gene-expression data have been deposited in Figshare (10.6084/m9.figshare.15152391)[59]. Data analysis scripts, implemented in Matlab, can be found at GitHub, https://github.com/sisichen-dev/NSC-Pop-Analysis. The scripts include a Matlab implementation of the PopAlign package[22], and requires several outside computing packages including the NMF toolbox[60], mmread (provided by NIST Matrix Market), and confplot[61].

## Acknowledgements

We would like to thank Allan Pool-Hermann, Eric Chow, Yaron Antebi, Zev Gartner, Michael Elowitz, David Brown, Pranav Bhamidipati, David Schaffer, Carlos Lois, Chris McGinnis, Jase Gehring, Elliott Robinson for feedback and discussions; Sandy Nandagopal, Elisha Mackay for reagents and cells; Inna-Marie Strahznik for figure illustrations, members of the Thomson Lab, and the Beckman Institute Single-cell Profiling and Engineering Center (SPEC). Funding support for this project was provided by the NIH Office of the Director (5DP5OD012194-04), the Shurl and Kay Curci Foundation, and the Heritage Medical Research Institute.

