## Supplementary material for "Designing signaling environments to steer transcriptional diversity in neural progenitor cell populations": SI

### Supplementary Methods

**1 Supplementary Methods** **3**

1.1 Data normalization . . . . . . . . . . . . . . . . . . . . . . . . . . . . 3

1.2 Probabilistic models and alignment . . . . . . . . . . . . . . . . . . . . 3

1.3 Query . . . . . . . . . . . . . . . . . . . . . . . . . . . . . . . . . . . 4

1.4 Statistical model of population response to signal concentrations . . . . . . . . . . 4

### Supplementary Figures

**S1 Building a GMM model from single-cell gene expression data.** . . . . . . . . . . 8

**S2 PCA plot of developmental timepoints, with mature cell type markers.** . . . . . 9

**S3 Model matches experimental data qualitatively.** . . . . . . . . . . . . . . . . . . 10

**S4 Quantitative comparison of model error for all samples.** . . . . . . . . . . . . . 11

**S5 Signaling pathways (BMP, PDGF, EGF, FGF, WNT, and Notch) are active in cells from the ventricular zone at E18.** . . . . . . . . . . . . . . . . . . . . . . . . 12

**S6 Gene expression heatmap of pre-astrocytic, pre-neuronal, and committed neuroblast cells from the combinatorial signaling screen.** . . . . . . . . . . . . . . . 13

**S7 Heatmaps of pre-astrocytic and committed neuroblast population proportions in response to additional signals.** . . . . . . . . . . . . . . . . . . . . . . . . . . 14

**S8 Ranking of samples relative to SVZ-D0 control.** . . . . . . . . . . . . . . . . . . . . 15

**S9 Coefficients comparison between the LS model and the log-linear model.** . . . 16

### Supplementary Table

**S1 All Dataset Information** . . . . . . . . . . . . . . . . . . . . . . . . . . . . . 17

**S2 Combinatorial Signaling Dataset Information.** All concentrations are given in ng/mL unless otherwise specified . . . . . . . . . . . . . . . . . . . . . . . . . . 18

### 1 Supplementary Methods

#### 1.1 Data normalization

Gene expression values are normalized according to the PopAlign paper[1]. Briefly, gene expression values, $g_i$, are first rescaled by the total number of transcripts per cell to account for droplet-to-droplet variation in transcript capture. Normalized values are then log-transformed after multiplication by a scaling factor $\beta$.

$$g_i' = \log(\beta \frac{g_i}{\sum_i^n g_i} + 1) \tag{1}$$

The scaling factor $\beta$ is chosen in order to achieve a smooth transition in the distribution of transformed $g_i'$ when raw $g_i$ values step from 0 to 1. We found that by setting $\beta$ to be roughly the median number of total transcript counts in a cell ( 3000) generally yields good results.

#### 1.2 Probabilistic models and alignment

To compare single-cell datasets, we construct probabilistic models that estimate the underlying joint probability distribution of cells within gene expression space. Probabilistic models are useful because they are a highly-compressed, conceptually meaningful representations of single-cell data, and their sub-components can be quantitatively compared using well-defined statistical metrics. In this work, the probabilistic model-based comparisons allow us to automatically align in vitro-grown cells to cells from the natural brain using statistical divergence, and rank and compare entire populations of cells using a log-likelihood ratio metric.

Models are constructed as described in the PopAlign paper [1]. Briefly, single-cell gene expression data is first normalized as described above. The genes are then filtered and only highly variable genes (e.g. supra-Poisson) are kept.

We then compress gene expression data into a set of m features, which are gene expression programs extracted from the data using matrix factorization techniques. In this work, we use the top 10- PCA features for analysis. Using these features, we construct a compressed representation of each cell as a vector $\boldsymbol{c} = (c_1, c_2, \ldots, c_m)$ of m feature coefficients, which weight the magnitude of gene expression programs within a given cell.

For each sample we build a mixture model that factors the population into a set of distinct subpopulations, each represented by a distinct mixture component (e.g. a Gaussian probability density):

$$P(\boldsymbol{c}) = \sum_{j=1}^{l} w_j \, \phi_j(\boldsymbol{c}) \tag{2}$$
$$\text{where } \phi_j(\boldsymbol{c}) = \mathcal{N}(\boldsymbol{c}; \boldsymbol{\mu_j}, \boldsymbol{\Sigma_j})$$

**Aligning probabilistic models** To determine which subpopulations to compare between samples, we perform an alignment between models based on statistical 'closeness'. We denote the population to be compared as the 'reference' population, and the other populations as 'test' populations. For each mixture component in the 'test' population, we can find the closest mixture component within the reference set by minimizing the Jeffreys divergence, a symmetrized version of the KL divergence:

$$\arg\min_j D_{\text{JD}} \left(\phi_i^{\text{test}}(\boldsymbol{c}) \parallel \phi_j^{\text{ref}}(\boldsymbol{c})\right), \tag{3}$$

#### 1.3 Query

In samples where the number of cells is too small to build a standalone model, we use an existing model to classify cells into different subpopulations. For the combinatorial signaling screen, some samples had roughly 100 cells, which would not adequately sample the feature space. For these experiments, we query each datapoint $c_i$ against a control model built from a subsample of the entire dataset. The control model consists of a collection of mixture components $\{\phi_j(\boldsymbol{c})\}$. We can then assign a class $Y_i$ for each specific cell - $\boldsymbol{c_i}$ :

$$Y_i = \arg\max_j \left(\phi_i(\boldsymbol{c_i})\right)$$

All cells in each sample receive a class label $Y$, these class labels can then be used to compute sample-specific subpopulation parameters - $\mu_j$, $\Sigma_j$ and $w_j$. The resulting proportional abundances $w_j$ are used in the following model to learn how population structure is steered by signals.

#### 1.4 Statistical model of population response to signal concentrations

Changing the signaling environment is a common strategy for the developing embryo to alter its population structure in accordance with developmental needs. Understanding how complex combinations of signals interact to regulate the relative abundance of populations within a complex mixture is critical for uncovering design principles that allow us to program populations of distinct stem and progenitor cell populations.

In our experimental system, we can apply many signaling concentrations on dissociated cells from the developing brain to understand how the populations are affected by single signals or their combinations. Many of the signals modulate the relative proportions of specific subpopulations within our heterogeneous mixture. For instance, we found that increasing EGF/FGF increases the proportion of pre-astrocytic progenitor cell type, while decreasing the proportion of the committed neuronal cell type.

We quantified the effects of each signaling molecules by a log-linear model that encodes growth rate of each cell type as a linear function of input signal concentrations. Prior work has shown that in individual cell types, combinations of signals can be integrated by cellular signal processing in potentially complex ways, including ratiometric detection [2], antagonism [3, 4, 5], or dominance [6]. However, additive responses to signaling combinations have been shown to one of the most common modes of signal integration [2], so we hypothesized that a linear model could be reasonably capture the population-level restructuring we observed in our signal-response data.

The model considers that each population $i$ undergoes independent exponential growth

$$n_i(t = T) = n_i(t = 0)\, e^{rT} \tag{4}$$

where $n_i$ is the cell number of population $i$, $r$ is the growth rate, and $T$ is the total growth time. We assume that the growth rate $r$ is modulated by the input signals by a linear function

$$r_i = \boldsymbol{a}_i^{\intercal}\boldsymbol{x} + b_i\,, \tag{5}$$

then the log of population is also a linear function of input signals

$$\log n_i(t = T) = T\boldsymbol{a}_i^{\intercal}\boldsymbol{x} + Tb_i + \log n_i(t = 0) \tag{6}$$

since $T$ and $n_i(t = 0)$ are constants. In the equation $\boldsymbol{x} = (x_1, x_2 \ldots, x_j)^{\intercal}$ are the concentrations of signaling molecules. For simplicity of notation, we denote the linear relationship as

$$\log n_i(t = T) = \boldsymbol{\beta_i^{\intercal}}\boldsymbol{x} + c_i\,. \tag{7}$$

We examined two models based on the linear relationship:

1. (LS model) Fitting the log cell counts with least squares
2. (Log-linear model) Fitting the ratio between cell types with a log-linear model

The fitting error of the LS model is rather large. The mean absolute error of log counts in the LS model is $0.459$, which means there is an average of $\exp 0.459 = 1.58$ fold difference in the model fitting and the data, as shown in SI Figure [add ref]. Nonetheless, the log-linear model gives a great fit to the proportions of cell types, with a mean absolute error $6.52\%$.

The results shows that the independent linear regulation of signaling cues generally hold, though in different experiments the total cell number could have experimental variance when taking samples from cell culture. Despite the different goodness-of-fit, the regulatory coefficients $\beta_i$s are generally consistent between the LS model and the log-linear model, as shown in SI Figure **??**. Therefore our analysis and interpretation of signaling effects are robust to the type of models.

Therefore, we only focus on the proportion of each cell population using the log-linear model in the following analysis. The proportion $p_i$ of population $i$ at time $T$ follows a log-linear distribution

$$p_i = \frac{n_i(t=T)}{\sum_j n_j(t=T)} = \frac{1}{\mathcal{Z}} \exp(\boldsymbol{\beta}_{\boldsymbol{i}}^\intercal \boldsymbol{x} + c_i)\,, \tag{8}$$

where the partition function $\mathcal{Z} = \sum_j \exp(\boldsymbol{\beta}_{\boldsymbol{j}}^\intercal \boldsymbol{x} + c_j)$. The coefficients $\beta_i$ of each population can be solved by maximum likelihood method.

The log likelihood function of $M$ different independent signaling condition is

$$\log \mathcal{L} = \log \prod_{m=1}^{M} \prod_i p_i^{(m)} = \sum_{m=1}^{M} \sum_i \log \frac{1}{\mathcal{Z}} \exp(\boldsymbol{\beta}_{\boldsymbol{i}}^\intercal \boldsymbol{x}^{(m)} + c_i)\,, \tag{9}$$

where $p_i^{(m)}$ is the proportion of population $i$ in $m$-th experiment. The parameters $\boldsymbol{\beta}_i$ and $c_i$ are determined by the optimization problem

$$\max_{\boldsymbol{\beta}_{\boldsymbol{i}}, c_i} \quad \log \mathcal{L}\,. \tag{10}$$

The maximum likelihood problem can be solved by typical gradient ascent. Also note that the current parameterization is not independent. Due to the constraints of probability summing to 1, the parameters of one cell type $k$ can be cancelled out in the probability expression

$$p_i = \frac{\exp(\boldsymbol{\beta}_{\boldsymbol{i}}^\intercal \boldsymbol{x} + c_i)}{\sum_j \exp(\boldsymbol{\beta}_{\boldsymbol{j}}^\intercal \boldsymbol{x} + c_j)} = \frac{\exp[(\boldsymbol{\beta}_{\boldsymbol{i}}^\intercal - \boldsymbol{\beta}_{\boldsymbol{k}}^\intercal)\boldsymbol{x} + (c_i - c_k)]}{\sum_j \exp[(\boldsymbol{\beta}_{\boldsymbol{j}}^\intercal - \boldsymbol{\beta}_{\boldsymbol{k}}^\intercal)\boldsymbol{x} + (c_j - c_k)]}\,. \tag{11}$$

Therefore we only able to determine the relative growth rate parameter $\boldsymbol{\beta}_i - \boldsymbol{\beta}_k$ and $c_i - c_k$. For the reference population $k$ the relative parameters $\boldsymbol{\beta}_k - \boldsymbol{\beta}_k = \mathbf{0}$, $c_k - c_k = 0$. For the simplicity of interpretation, the reference population $k$ should be relatively stable to signals. Thus we choose cell type 3 (pre-neuronal progenitors) as the reference given that its proportions are the most consistent across all conditions.

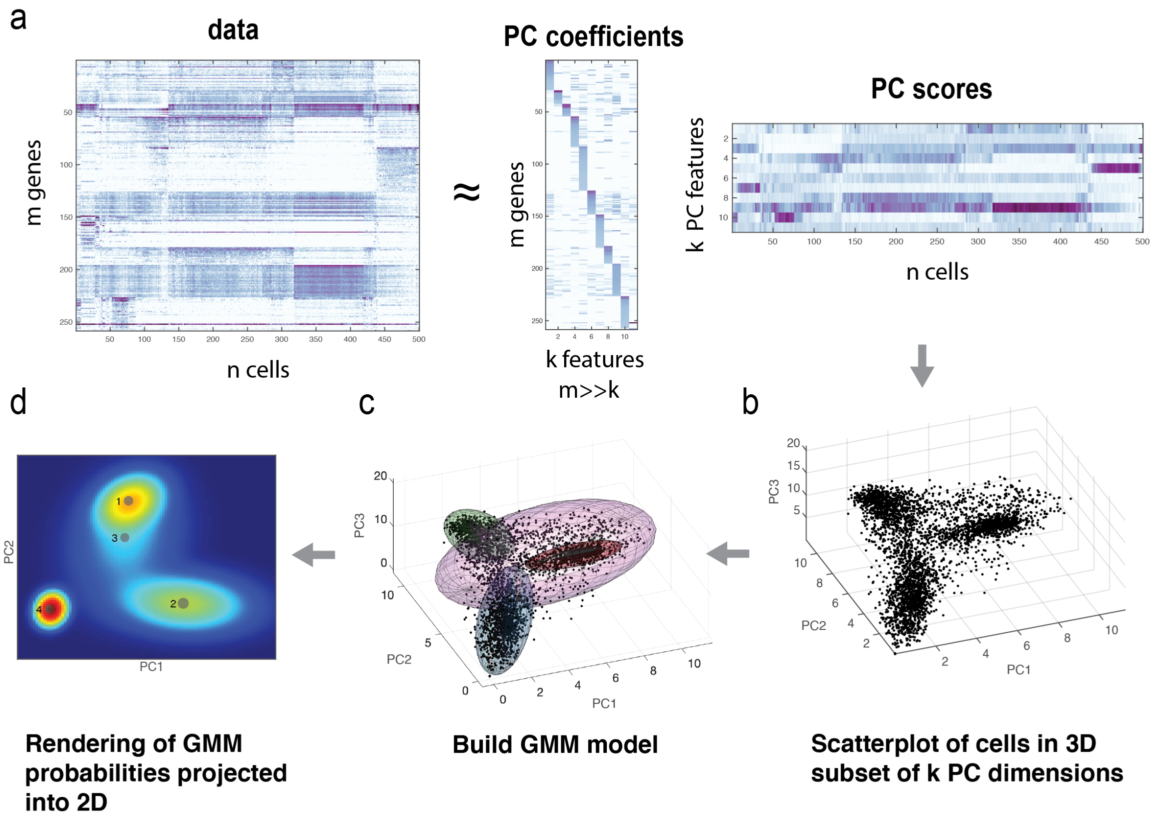


**SI Figure S1: Building a GMM model from single-cell gene expression data.** a) Normalized, and filtered single-cell gene expression data matrix is dimensionality reduced using PCA. PC coefficient vectors are truncated to k dimensions (k=10 in our analysis). (b) Scatter plot showing cells in a 3-d subset of the principal component dimensions. (c) GMM model is built within the k-dimensional space (3d subset shown in figure) (d) Rendering of GMM predicted probabilities projected down onto first two PCs.

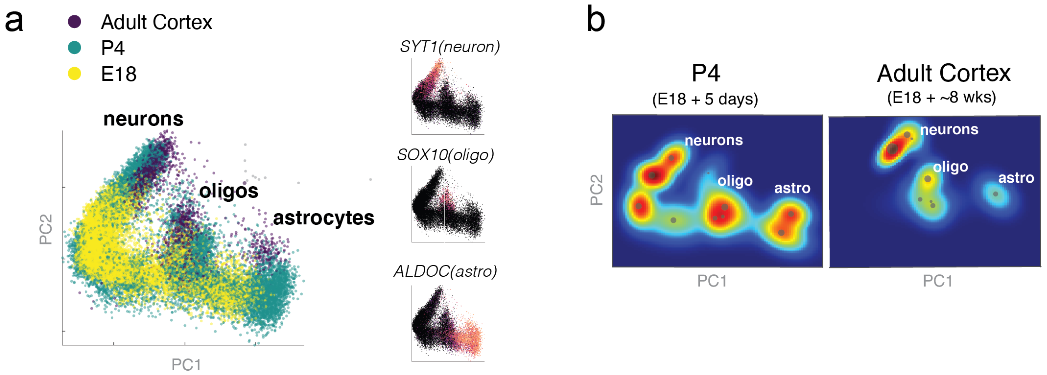


**SI Figure S2: PCA plot of developmental timepoints, with mature cell type markers.** (a) Scatterplots of single-cells from all timepoints in PC space, colored by developmental stage. Inset plots show expression patterns of mature neuron marker (SYT1), mature oligodendrocyte marker (SOX10) and mature astrocyte marker (ALDOC). (b) Heatmap rendering of GMM models for timepoints P4 and adult cortex, with regions containing differentiated cell types labeled according to (a)

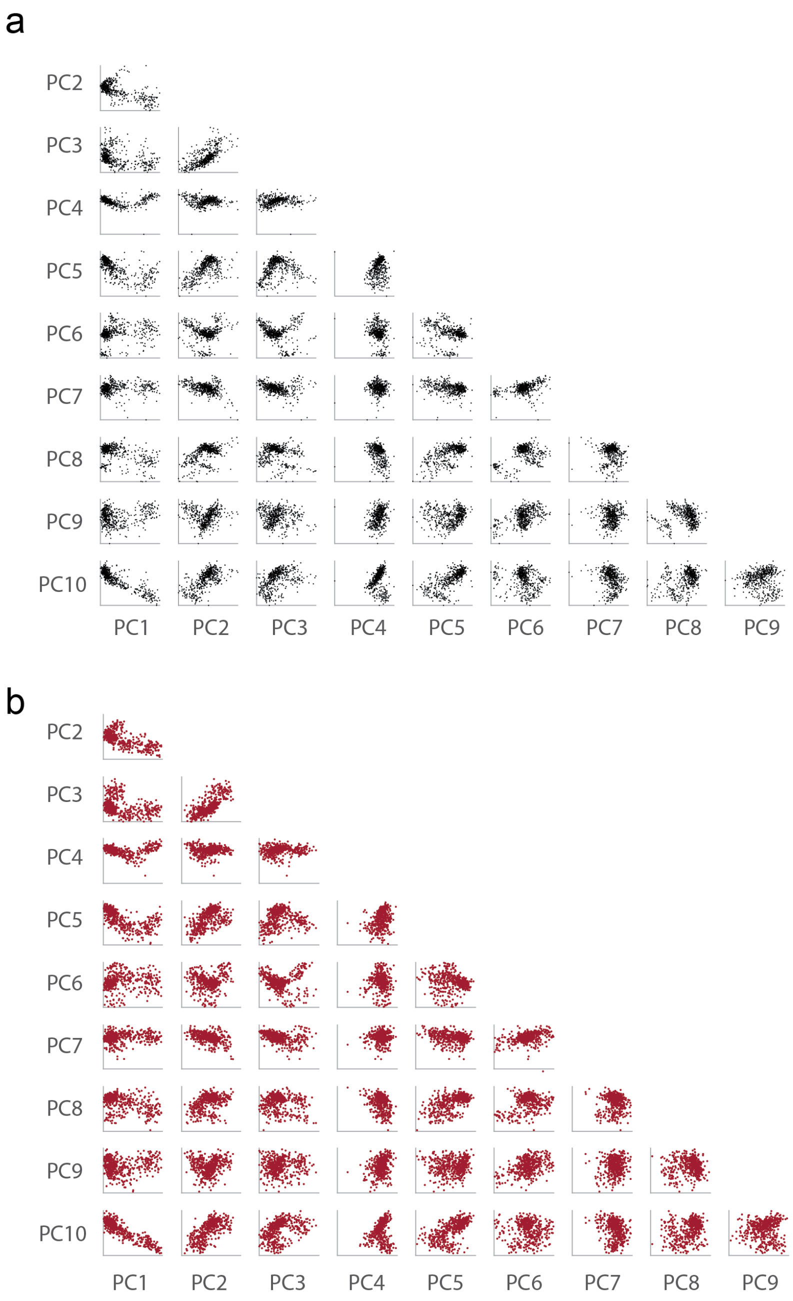


**SI Figure S3: Model matches experimental data qualitatively.** Array of 2D scatterplots of experimental data for the (a) SVZ - D0 sample and (b) model-simulated data. For each plot tile, the PC dimension is are displayed on the axis. Corresponding plots have the same x- and y- axis scaling.

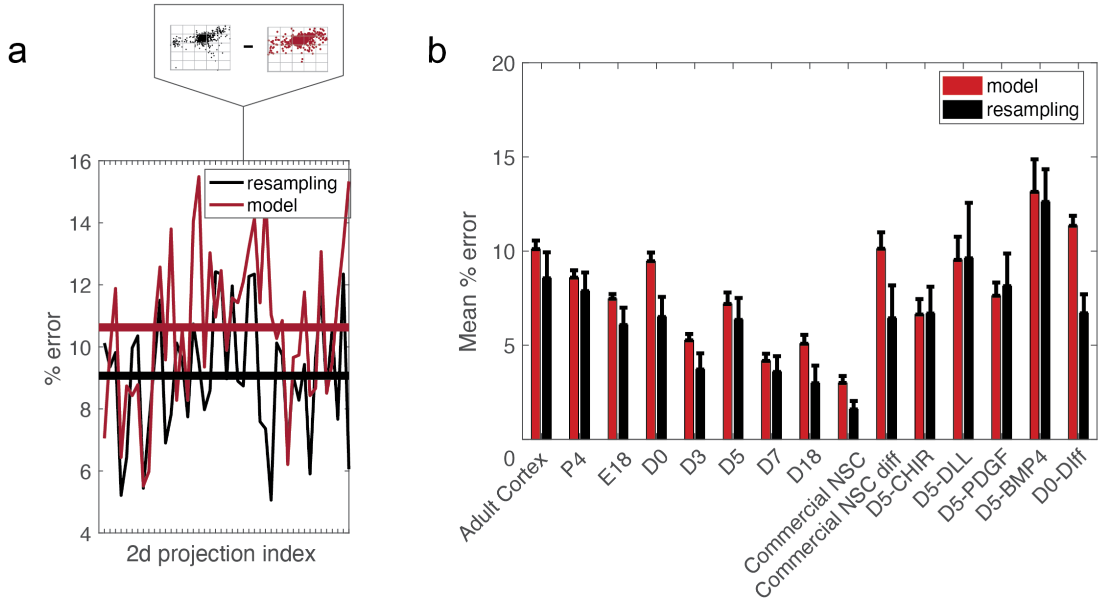


**SI Figure S4: Quantitative comparison of model error for all samples.** (a) % Error for each 2D projection, calculated across a 5x5 array of 2D bins (see Methods). Model error is in red and sampling error (calculated between training and validation data) is in black. Mean % error across all 2D projections is denoted as thick horizontal line. (b) Mean % error across all 2D projections is compared to data sampling error for each dataset. Error bars show the standard deviation calculated across 20 separate model building and data resampling attempts.

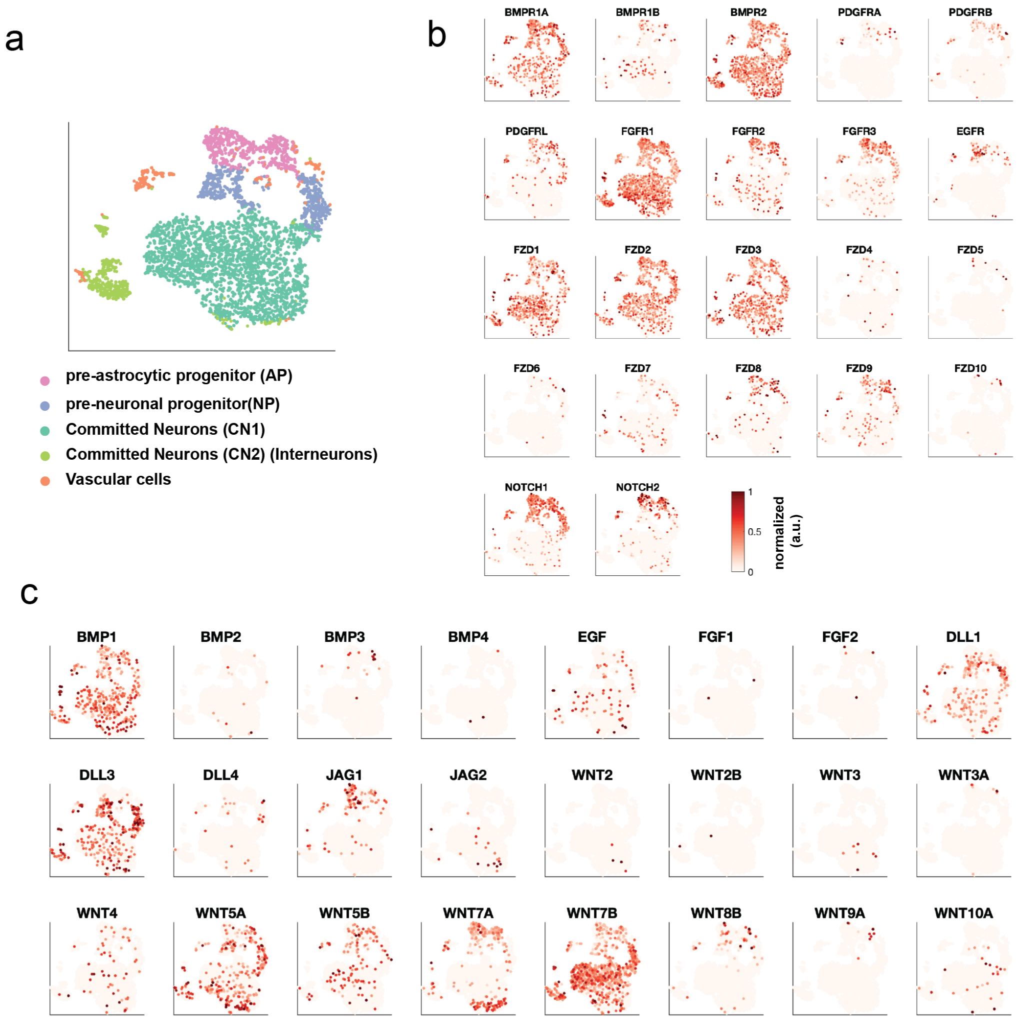


**SI Figure S5: Signaling pathways (BMP, PDGF, EGF, FGF, WNT, and Notch) are active in cells from the ventricular zone at E18.** (a) t-SNE plot of VZ-D0 cells separated into distinct subpopulations. (b) Normalized gene expression values for receptors of the aforementioned signaling pathways. (c) Normalized gene expression values for ligands of the aforementioned signaling pathways. All data are log-normalized and rescaled from 0 to 1.

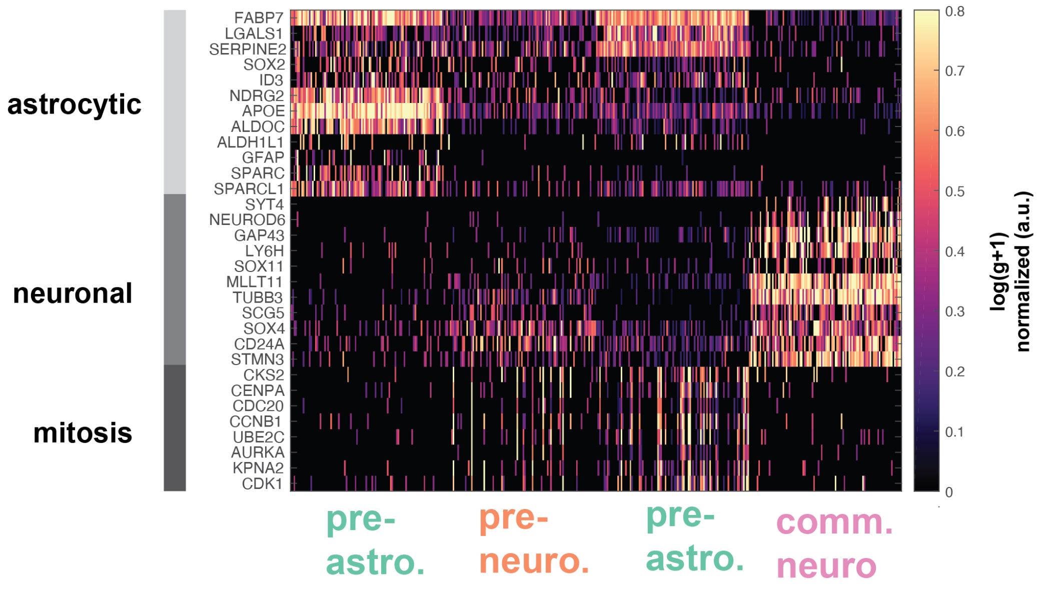


**SI Figure S6: Gene expression heatmap of pre-astrocytic, pre-neuronal, and committed neuroblast cells from the combinatorial signaling screen.** Gene expression programs are denoted on the right. Gene counts are normalized and logged (see Methods), and then rescaled by the gene maximum in each row. Displayed values are saturated at 0.8 to improve visual clarity.

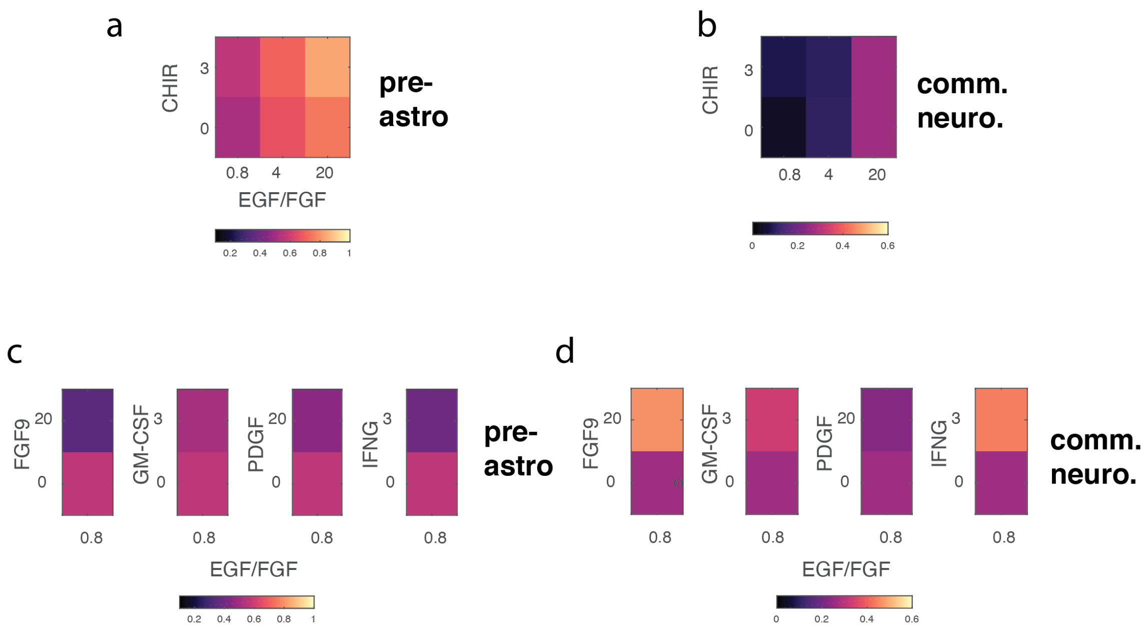


**SI Figure S7: Heatmaps of pre-astrocytic and committed neuroblast population proportions in response to additional signals.** The proportion of (a) pre-astrocytic population and (b) committed neuroblast populations in response to CHIR and EGF/FGF. The proportions of (c) pre-astrocytic and (d) committed neuroblastpopulations in response to FGF9, GM-CSF, PDGF, IFN-G, at low EGF/FGF concentrations (0.8 ng/mL).

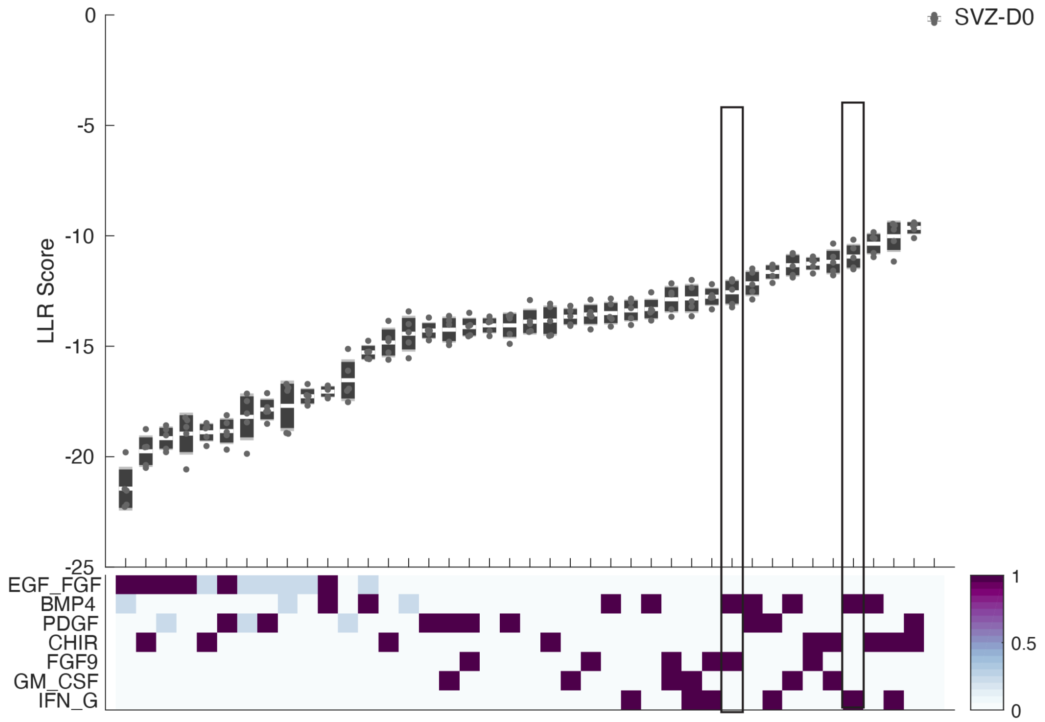


**SI Figure S8: Ranking of samples relative to SVZ-D0 control.** (a) Log-likelihood ratio (LLR) scores for each sample against the control population. The more a population of cells diverges from the SVZ-D0 sample, the lower its LLR score. The two conditions designed in Fig 4e-f are highlighed in boxes. LLR scores are bootstrapped by building 20 models for each signaling ondition and then calculating the LLR relative to the SVZ-D0 model, and using 100 cells are sampled from each signaling condition. White line: mean. Dark gray: SEM. Light gray: std. Outliers are individually plotted.

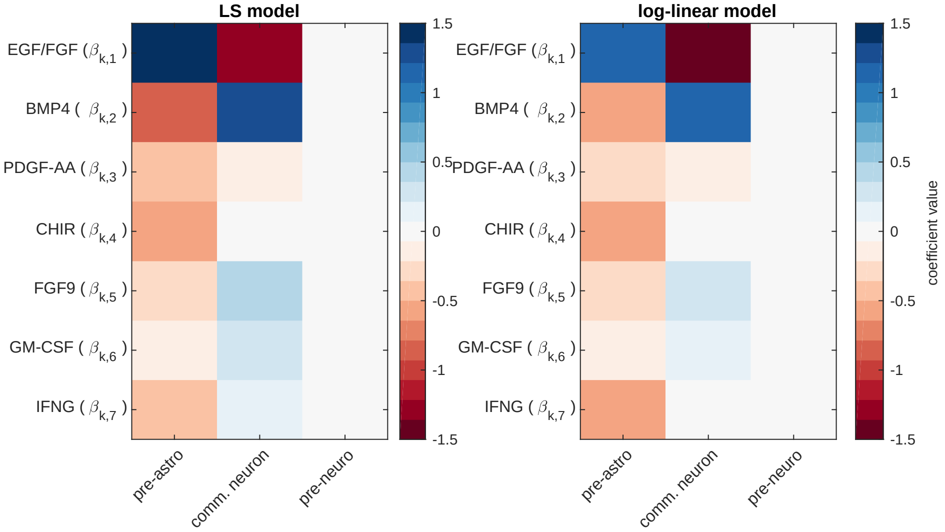


**SI Figure S9: Coefficients comparison between the LS model and the log-linear model.**

**SI Table S1: All Dataset Information**

| Name | Description | # cells | Type |
|---|---|---|---|
| P4 | Whole brain from P4 mice | 12015 | live |
| AdCo | Cortex from 8-week old mouse | 2657 | live |
| E18 (10x Genomics) | combined cortex, hippocampus, and ventricular zone | 20000 | live |
| SVZ-D0 | E18 combined cortex, hippocampus, and ventricular zone | 5030 | live |
| SVZ-D3 | 3 days in Npgrow | 9862 | live |
| SVZ-D5 | 5 days in Npgrow | 1125 | live |
| SVZ-D7 | 7 days in Npgrow | 3820 | live |
| SVZ-D18 | 18 days in Npgrow | 6256 | live |
| SVZ-D0-DIFF | 5 days in NBco-culture media | 3679 | live |
| NPC-D5-GROWTH | 5 days in NPgrow | 10527 | live |
| NPC-D5-DIFF | 5 days in NPgrow, then 5 days in co-culture media | 3537 | live |
| SVZ-D5-CHIR | 5 days in NPgrow + CHIR (3 uM) | 997 | live |
| SVZ-D5-PDGF | 5 days in NPgrow + PDGF (20ng/mL) | 903 | live |
| SVZ-D5-BMP4 | 5 days in NPgrow + BMP4 (20ng/mL) | 715 | live |
| NPC-MULT-1 | 5 days in NPgrow (minus EGF/FGF) + supplemental factors | 8691 | fixed |
| NPC-MULT-2 | 5 days in NPgrow (minus EGF/FGF) + supplemental factors | 10020 | fixed |

**SI Table S2: Combinatorial Signaling Dataset Information.**
All concentrations are given in ng/mL unless otherwise specified

| sample_name | dataset | EGF_FGF | BMP4 | PDGF | CHIR(uM) | FGF9 | GM_CSF | IFN_G |
|---|---|---|---|---|---|---|---|---|
| EGF_FGF-20 | NPC-MULT-1 | 20 | 0 | 0 | 0 | 0 | 0 | 0 |
| EGF_FGF-4 | NPC-MULT-1 | 4 | 0 | 0 | 0 | 0 | 0 | 0 |
| EGF_FGF-0.8 | NPC-MULT-1 | 0.8 | 0 | 0 | 0 | 0 | 0 | 0 |
| EGF_FGF-20_BMP4-4 | NPC-MULT-1 | 20 | 4 | 0 | 0 | 0 | 0 | 0 |
| EGF_FGF-4_BMP4-4 | NPC-MULT-1 | 4 | 4 | 0 | 0 | 0 | 0 | 0 |
| EGF_FGF-0.8_BMP4-4 | NPC-MULT-1 | 0.8 | 4 | 0 | 0 | 0 | 0 | 0 |
| EGF_FGF-20_BMP4-20 | NPC-MULT-1 | 20 | 20 | 0 | 0 | 0 | 0 | 0 |
| EGF_FGF-4_BMP4-20 | NPC-MULT-1 | 4 | 20 | 0 | 0 | 0 | 0 | 0 |
| EGF_FGF-0.8_BMP4-20 | NPC-MULT-1 | 0.8 | 20 | 0 | 0 | 0 | 0 | 0 |
| EGF_FGF-20_PDGF-20 | NPC-MULT-1 | 20 | 0 | 20 | 0 | 0 | 0 | 0 |
| EGF_FGF-4_PDGF-20 | NPC-MULT-1 | 4 | 0 | 20 | 0 | 0 | 0 | 0 |
| EGF_FGF-0.8_PDGF-20 | NPC-MULT-1 | 0.8 | 0 | 20 | 0 | 0 | 0 | 0 |
| EGF_FGF-20_PDGF-4 | NPC-MULT-1 | 20 | 0 | 4 | 0 | 0 | 0 | 0 |
| EGF_FGF-4_PDGF-4 | NPC-MULT-1 | 4 | 0 | 4 | 0 | 0 | 0 | 0 |
| EGF_FGF-0.8_PDGF-4 | NPC-MULT-1 | 0.8 | 0 | 4 | 0 | 0 | 0 | 0 |
| EGF_FGF-20_CHIR-3 | NPC-MULT-1 | 20 | 0 | 0 | 3 | 0 | 0 | 0 |
| EGF_FGF-4_CHIR-3 | NPC-MULT-1 | 4 | 0 | 0 | 3 | 0 | 0 | 0 |
| EGF_FGF-0.8_CHIR-3 | NPC-MULT-1 | 0.8 | 0 | 0 | 3 | 0 | 0 | 0 |
| EGF_FGF-0.8_IFN_G-3 | NPC-MULT-2 | 0.8 | 0 | 0 | 0 | 0 | 0 | 3 |
| EGF_FGF-0.8_IFN_G-3_GM_CSF-3 | NPC-MULT-2 | 0.8 | 0 | 0 | 0 | 0 | 3 | 3 |
| EGF_FGF-0.8_IFN_G-3_PDGF-20 | NPC-MULT-2 | 0.8 | 0 | 20 | 0 | 0 | 0 | 3 |
| EGF_FGF-0.8_IFN_G-3_BMP4-20 | NPC-MULT-2 | 0.8 | 20 | 0 | 0 | 0 | 0 | 3 |
| EGF_FGF-0.8_IFN_G-3_FGF9-20 | NPC-MULT-2 | 0.8 | 0 | 0 | 0 | 20 | 0 | 3 |
| EGF_FGF-0.8_IFN_G-3_CHIR-3 | NPC-MULT-2 | 0.8 | 0 | 0 | 3 | 0 | 0 | 3 |
| EGF_FGF-0.8_FGF9-20 | NPC-MULT-2 | 0.8 | 0 | 0 | 0 | 20 | 0 | 0 |
| EGF_FGF-0.8_GM_CSF-3 | NPC-MULT-2 | 0.8 | 0 | 0 | 0 | 0 | 3 | 0 |
| EGF_FGF-0.8_GM_CSF-3_PDGF-20 | NPC-MULT-2 | 0.8 | 0 | 20 | 0 | 0 | 3 | 0 |
| EGF_FGF-0.8_GM_CSF-3_BMP4-20 | NPC-MULT-2 | 0.8 | 20 | 0 | 0 | 0 | 3 | 0 |
| EGF_FGF-0.8_GM_CSF-3_FGF9-20 | NPC-MULT-2 | 0.8 | 0 | 0 | 0 | 20 | 3 | 0 |
| EGF_FGF-0.8_GM_CSF-3_CHIR-3 | NPC-MULT-2 | 0.8 | 0 | 0 | 3 | 0 | 3 | 0 |
| EGF_FGF-0.8_FGF9-20_CHIR-3 | NPC-MULT-2 | 0.8 | 0 | 0 | 3 | 20 | 0 | 0 |
| EGF_FGF-0.8_CHIR-3 | NPC-MULT-2 | 0.8 | 0 | 0 | 3 | 0 | 0 | 0 |
| EGF_FGF-0.8_PDGF-20 | NPC-MULT-2 | 0.8 | 0 | 20 | 0 | 0 | 0 | 0 |
| EGF_FGF-0.8_PDGF-20_BMP4-20 | NPC-MULT-2 | 0.8 | 20 | 20 | 0 | 0 | 0 | 0 |
| EGF_FGF-0.8_PDGF-20_FGF9-20 | NPC-MULT-2 | 0.8 | 0 | 20 | 0 | 20 | 0 | 0 |
| EGF_FGF-0.8_PDGF-20_CHIR-3 | NPC-MULT-2 | 0.8 | 0 | 20 | 3 | 0 | 0 | 0 |
| EGF_FGF-0.8 | NPC-MULT-2 | 0.8 | 0 | 0 | 0 | 0 | 0 | 0 |
| EGF_FGF-0.8_BMP4-20 | NPC-MULT-2 | 0.8 | 20 | 0 | 0 | 0 | 0 | 0 |
| EGF_FGF-0.8_BMP4-20_FGF9-20 | NPC-MULT-2 | 0.8 | 20 | 0 | 0 | 20 | 0 | 0 |
| EGF_FGF-0.8_BMP4-20_CHIR-3 | NPC-MULT-2 | 0.8 | 20 | 0 | 3 | 0 | 0 | 0 |
